# Advancing childhood cancer research through young investigator and advocate collaboration

**DOI:** 10.1101/2023.12.03.569769

**Authors:** Amber K. Weiner, Antonia Palmer, Melanie Frost Moll, Gavin Lindberg, Kevin Reidy, Sharon J. Diskin, Crystal L. Mackall, John M. Maris, Patrick J. Sullivan

**Author notes:** Correspondence: John M. Maris - 3401 Civic Center Blvd, Philadelphia, PA 19104.

## Abstract

Cancer advocates and researchers share the same goal of driving science forward to create new therapies to cure more patients. The power of combining cancer researchers and advocates has become of increased importance due to their complementary expertise. Therefore, advocacy is a critical component of grant structures and has become embedded into the Stand Up 2 Cancer (SU2C) applications. To date, the optimal way to combine these skillsets and experiences to benefit the cancer community is currently unknown. The Saint Baldrick’s Foundation (SBF)–SU2C now called St. Baldrick’s Empowering Pediatric Immunotherapies for Childhood Cancer (EPICC) Team is comprised of a collaborative network across nine institutions in the United States and Canada. Since SU2C encourages incorporating advocacy into the team structure, we have assembled a diverse team of advocates and scientists by nominating a young investigator (YI) and advocate from each site. In order to further bridge this interaction beyond virtual monthly and yearly in person meetings, we have developed a questionnaire and conducted interviews. The questionnaire is focused on understanding each member’s experience at the intersection between science/advocacy, comparing to previous experiences, providing advice on incorporating advocacy into team science and discussing how we can build on our work. Through creating a YI and advocate infrastructure, we have cultivated a supportive environment for meaningful conversation that impacts the entire research team. We see this as a model for team science by combining expertise to drive innovation forward and positively impact pediatric cancer patients, and perhaps those with adult malignancies.

**Significance:** Questionnaire results show both advocates and YI’s see this structure to be valuable and beneficial. YI’s communicated their research to a non-scientific audience and learned advocate’s experience. This was their first advocacy experience for most YIs. Advocates learned more about the research being conducted to provide hope. They can also aid with fundraising, publicity and lobbying. This collaboration improves science communication, designing patient-friendly clinical trials and sharing experience across institutions.

## Introduction

Cancer researchers and advocates share the common goal of driving science forward to create new therapies and cure more patients [1-2]. Both constituencies have the prospect of accelerating the realization of that common goal by combining their shared passions and complementary expertise [3]. The optimal way to combine these skill sets and experiences to benefit the cancer community is currently unknown.

The St. Baldrick’s Foundation (SBF)-Stand Up 2 Cancer (SU2C) Pediatric Cancer Dream Team (PCDT) is comprised of a highly collaborative network across the United States and Canada. Advocacy is a critical component of the SBF-SU2C program since its inception in 2013 and is embedded into all SU2C grant applications, requiring the incorporation of advocacy into the team structure.

Initially, the PCDT was formed by investigators with complementary expertise across seven institutions and, as required by the grant structure, six patient advocates (**Figure 1**). There was enthusiasm to incorporate advocacy into the team but there was a lack of experience and guidelines on the best approach.

**Figure 1.**
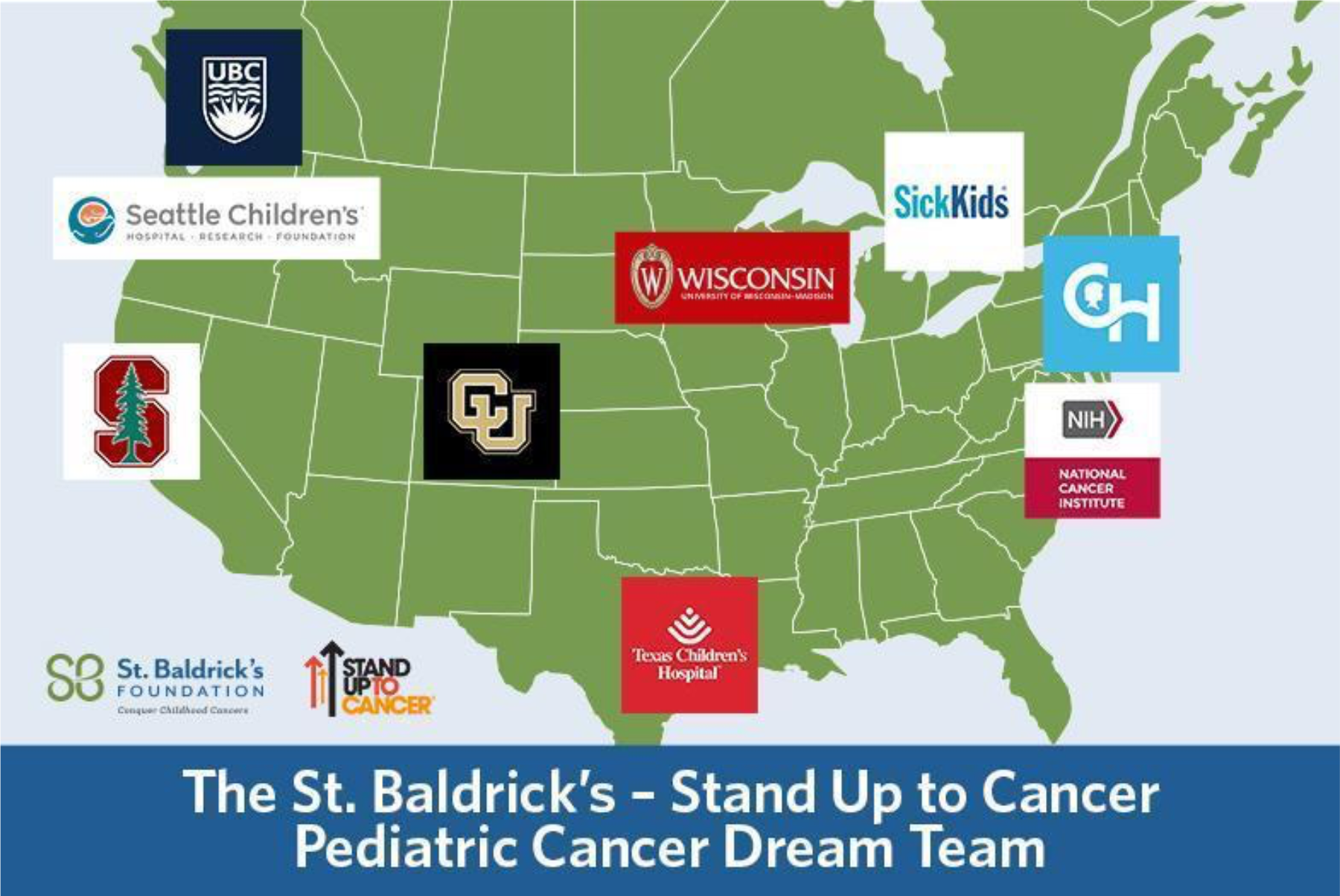
Institutions collaborating on the St. Baldrick Foundation (SBF)-SU2C Pediatric Cancer Dream Team (PCDT). The SBF-SU2C PCDT is comprised of a highly collaborative network across nine institutions in the United States and Canada. Since SU2C encourages incorporating advocacy into the team structure, we have assembled a diverse team of advocates and scientists by nominating a YI (YI) and advocate from each site. Image obtained from: https://www.stbaldricks.org/blog/post/pediatric-cancer-dream-team-works-toward-more-breakthroughs

During the second four years of funding, there was a strategic planning committee formed to outline future plans and directions with a specific focus on how better to incorporate the particular skill sets of advocates to advance the objectives of the team. A decision was made to strategically focus on enhancing interactions between advocates and young investigators (YIs) with the goal of teaching YIs how to effectively communicate their science and for advocates to learn more about the scientific underpinning of the team.

To pursue this strategic objective, an advocate and YI was nominated at each of the participating institutions to work together as part of the network. To build on this and establish a meaningful partnership, a pre-meeting of advocates and YIs was held before the 2018 annual meeting. This pre-meeting was dedicated to the education of both advocates and YIs and fostering the relationships between advocates and YIs across institutions. In 2019, the advocates and YIs hosted “A Night of Inspiration” where advocates and YIs from each site jointly shared our stories and research progress.

In 2020, a joint virtual training session was held with the advocates, YIs, and Jon Retzlaff, Chief Policy Officer at the American Association for Cancer Research (AACR). Monthly, the advocates have calls that often include a YI to present their research in lay terms, moderated by the senior investigator team leaders. Last year advocates collaboratively designed the website and created a logo for the PCDT (**Figure 2**). Since 2022, The PCDT is now known as the St. Baldrick’s Empowering Pediatric Immunotherapies for Childhood Cancer (EPICC) Team (PCDTeam.org).

**Figure 2.**
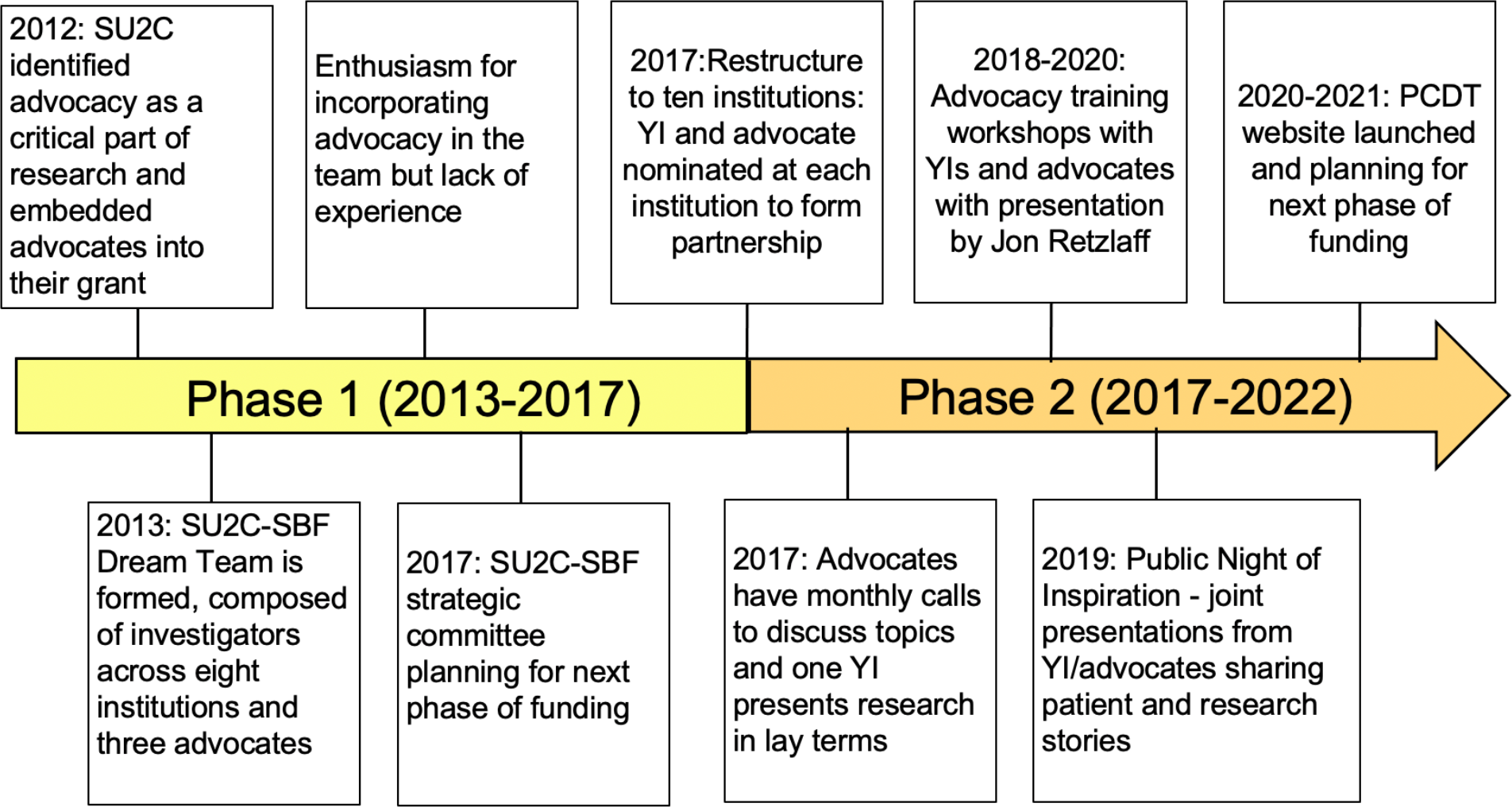
Timeline of Pediatric Cancer Dream Team Phases. The timeline highlights the milestones achieved during the two phases of funding.

In order to understand the impact of our efforts, we developed a questionnaire and conducted interviews with advocates and YIs. The questionnaire focused on understanding each member’s experience at the intersection between science and advocacy, comparing to previous experiences, providing advice on incorporating advocacy into team science, and discussing how to build on our work. We analyzed our interview data using grounded theory [4] where collected data was iteratively coded, categorized, and multiple “theories” identified. Our study identified several themes centered on communication, establishing meaningful and beneficial partnerships, the importance of early involvement of the advocates, and a joint focus on philanthropic fundraising, lobbying, and traditional grant funding.

Overall, the SBF-SU2C PCDT now known as the St. Baldrick’s EPICC Team provided the opportunity to incorporate advocacy into team science, which resulted in enriching research and promoting new and creative ways of thinking. Our infrastructure was developed by the advocates in collaboration with leadership, which resulted in dual learning objectives to influence the culture of YIs. Our analysis has demonstrated the importance and value of incorporating advocates with YIs in team science. This systemic interaction permits YIs to practice scientific communication and advocates to learn science terminology. Overall, the results demonstrate that integrating YIs and advocates in the context of a large team grant is a meaningfully important mechanism of effecting a cultural shift where research is done *with* patients and not just *for* patients.

## Materials and Methods

### Questionnaire

The following questions were provided prior to interviews. YI and advocates discussed these questions during group interviews or provided individual written responses.

1. How did you become involved in advocacy/science intersection?
2. Were you involved at your home institution or have other experiences?
3. Was your involvement independent of SB-SU2C or because it was suggested?
4. If you had previous experience before this team, how did it compare?
5. Do they find the integration valuable (YI and Advocates) important?
6. How has your collaboration with the advocates informed the way you communicate your research to a non-scientific audience? (YI Only)
7. How can this interaction evolve and improve in the future?
8. Do you have advice to give others on the challenges of bringing advocates and scientists together?

### Interview Structure

We interviewed ten young investigators and eight advocates with five advocates providing written responses. The interview questions focused on understanding each member’s experience at the intersection between science and advocacy, comparing to previous experiences, providing advice on incorporating advocacy into team science, and discussing how we can build on our work. Interviews were conducted in groups of advocates and YIs from different institutions. Interviews were recorded with participant’s permission by video conferencing for subsequent grounded theory analysis.

### Grounded Theory Analysis

Interviews that were recorded with consent were then transcribed and coded using a grounded theory framework [4]. The grounded theory process was conducted in multiple iterations to ensure alignment on coded terms across all the interviews that were conducted through in-person meetings and written responses. Upon completion, code words were tabulated, which resulted in the phrases used for word cloud generation. Next, terms were grouped into categories, which resulted in our overall theories or themes.

## Results

### Qualitative Analysis and Themes

The grounded theory analysis of the interview data yielded 104 terms of interest. After counting the number of times the words appeared across all interviews, we filtered for those that were noted more than ten times. This yielded eighteen terms that were visualized using a word cloud (**Figure 3**)

**Figure 3.**
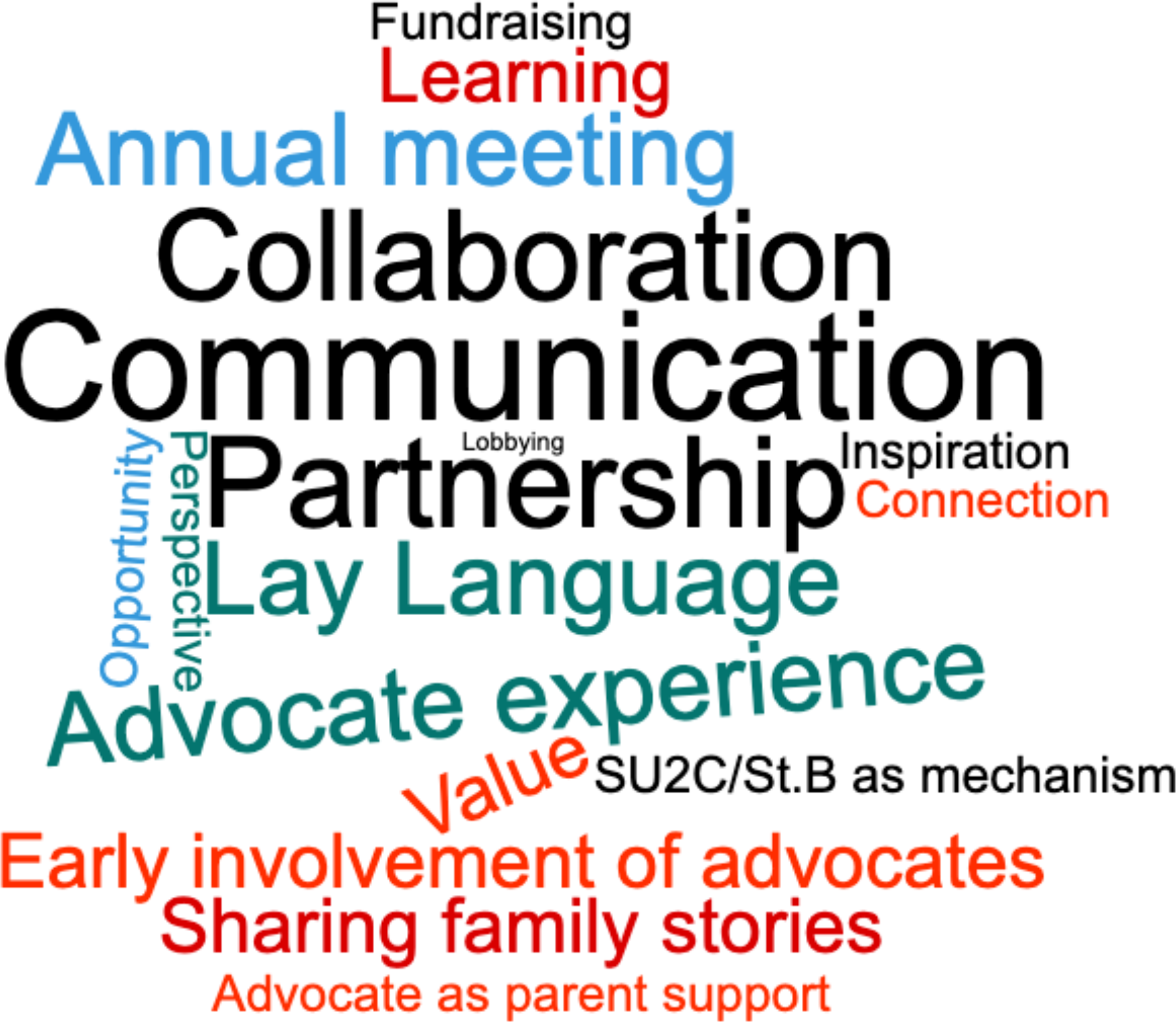
Word cloud of terms from grounded theory analysis of interviews. Interviews were transcribed and coded using a grounded theory analysis method. After multiple iterations, code words were tabulated, and data was used as word cloud input. The top 18 terms were used for word cloud generation.

Code words were grouped into theories or themes to draw overall conclusions. The top themes were: (1) **<u>Communicating</u>** regularly in lay language is essential for learning, sharing stories, providing a comfortable environment to build strong relationships, and maintaining momentum to reach goals. (2) YI and advocate **<u>partnership</u>** provides a unique opportunity to **<u>collaborate</u>** toward a common goal. SBF-SU2C PCDT **<u>annual meeting</u>** is one mechanism to connect as a team, maintain involvement and share inspiration. (3) The **<u>early involvement of advocates</u>** in the design of clinical trials is critical to address issues of quality of life, burden of care, patient accrual and retention. (4) **<u>Fundraising, lobbying, and funding</u>** are essential to change the environment, raise awareness and influence the next generation (5) YI and advocate partnership adds **<u>meaning and beneficial value</u>** by moving science forward and promoting career development to advance innovative therapies.

During the interview process, YIs and advocates provided many insights on the benefits of this relationship and how it creates unique and genuine learning experiences for all involved. One YI interviewee noted “This gives an opportunity for real dialogue where YIs get exposed early in their career before they have made many, many mistakes. Hopefully through some of these interactions we are making the road a little easier for the next set of investigators who come along.” Many YIs had never worked with advocates before in a formalized relationship as that experienced through the SBF-SU2C PCDT. One YI noted, “I can’t imagine any experience, also outside of the Dream Team, not involving advocates.” YIs and advocates both see this experience as being one that is valuable, lasting, and extending into other areas of their work.

## Discussion

All interviewees praised the design and implementation of the YI and advocate partnership, expressing its many benefits and value, despite being a significant investment in time. One young investigator spoke highly of the YI and advocate relationship, and in particular the “Night of Inspiration”, sharing that it was a way to “get this story out in a way that’s very tangible that’s very connected to both the science and the non-science groups and I think that’s the power of this group that’s unique. I think it should be expanded upon and used extensively because I don’t think that happens in very many scenarios and so I think that is a super weapon in my opinion”.

Several recommendations also resulted from the analysis and identified themes. (1) YIs and advocates all want regular contact and have the ability to meet on a more frequent basis as a larger group rather than once a year, (2) YIs and advocates would like the opportunity to support each other outside of their home institution (e.g., the advocate from one institution supporting a YI from a different institution), and (3) create more opportunities for training sessions that span multiple topics from science to advocacy.

To maintain momentum between yearly in-person meetings and bridge interactions between institutions, we have added monthly meetings focused on topics of interest. These meetings are an opportunity to provide training in science communication, legislation/government, regulatory science, and fundraising. We think that our collaborative partnership is a model for team science by combining expertise to drive innovation forward and positively impact pediatric cancer patients and is applicable to other patient populations with focused research programs.

We have formed a collaborative network of YIs and advocates committed to pediatric cancer translational immuno-oncology from institutions across North America. Through creating a YI and advocate infrastructure, we have cultivated a supportive environment for meaningful conversation that impacts the entire research team.

## Acknowledgements

We would like to thank all the study participants, which include the YIs (Kristopher Bosse, Christian Capitini, Sabina Kaczanowska, Robbie Majzner, Sneha Ramakrishna, Vijay Ramaswamy, Khaled Sanber, Corrine Summers, Amanda Winters, Haifeng Zhang) and patient advocates (Kelly Cotter, Kelly Forebaugh, Bambi Grilley, Gavin Lindberg, Carlos Sandi, Lori Schultz) on the St. Baldrick’s Foundation and Stand Up 2 Cancer Pediatric Cancer Dream Team. We would also like to acknowledge the nine institutions that are collaborating on this team.

The SBF-SU2C PCDT is led by John Maris and Crystal Mackall. The idea driving this work was developed in collaboration with them and Patrick Sullivan. All authors contributed to the study design and questionnaire. Amber Weiner organized all interviews with Antonia Palmer, Kevin Reidy and Melanie Moll participating in conducting the interviews. Semantic analysis was completed by Amber Weiner, Antonia Palmer and Melanie Moll. Figures and text were generated by Amber Weiner. All authors contributed to manuscript revision and editing.

This work was supported by a St. Baldrick’s-Stand Up To Cancer Pediatric Dream Team Translational Research Grant (SU2C-AACR-DT2727). Stand Up to Cancer is a program of the Entertainment Industry Foundation administered by the American Association for Cancer Research (JMM and CLM). This work was also supported by NCI R35 CA220500 (JMM).

